# Overexpressing *CsGH3.1* and *CsGH3.1L* reduces susceptibility to *Xanthomonas citri* subsp. *citri* by repressing auxin signaling in citrus (*Citrus sinensis* Osbeck)

**DOI:** 10.1101/697060

**Authors:** Xiuping Zou, Junhong Long, Ke Zhao, Aihong Peng, Min Chen, Qin Long, Yongrui He, Shanchun Chen

## Abstract

The auxin early response gene Gretchen Hagen3 (*GH3*) plays dual roles in plant development and responses to biotic or abiotic stress. It functions in regulating hormone homeostasis through the conjugation of free auxin to amino acids. In citrus, *GH3.1* and *GH3.1L* play important roles in responding to *Xanthomonas citri* subsp. *citri* (Xcc). Here, in Wanjingcheng orange (*C. sinensis* Osbeck), the overexpression of *CsGH3.1* and *CsGH3.1L* caused increased branching and drooping dwarfism, as well as smaller, thinner and upward curling leaves compared with wild-type. Hormone determinations showed that overexpressing *CsGH3.1* and *CsGH3.1L* decreased the free auxin contents and accelerated the Xcc-induced decline of free auxin levels in transgenic plants. A resistance analysis showed that transgenic plants had reduced susceptibility to citrus canker, and a transcriptomic analysis revealed that hormone signal transduction-related pathways were significantly affected by the overexpression of *CsGH3.1* and *CsGH3.1L*. A MapMan analysis further showed that overexpressing either of these two genes significantly downregulated the expression levels of the annotated auxin/indole-3-acetic acid family genes and significantly upregulated biotic stress-related functions and pathways. Salicylic acid, jasmonic acid, abscisic acid, ethylene and zeatin levels in transgenic plants displayed obvious changes compared with wild-type. In particular, the salicylic acid and ethylene levels involved in plant resistance responses markedly increased in transgenic plants. Thus, the overexpression of *CsGH3.1* and *CsGH3.1L* reduces plant susceptibility to citrus canker by repressing auxin signaling and enhancing defense responses. Our study demonstrates auxin homeostasis’ potential in engineering disease resistance in citrus.

## Introduction

Citrus canker, caused by *Xanthomonas citri* subsp. *citri* (Xcc), is an important disease of citrus. Xcc affects various citrus species and most of the economically important cultivars, including orange, grapefruit, lime, lemon, pomelo and citrus rootstock ^[1]^. The canker’s development includes the initial appearance of oily looking spots, usually on the abaxial leaf surface, outbursts of white or yellow spongy pustules and finally the formation of brown corky cankers ^[2]^. Pustule formation (excessive cell division) in the infected tissues plays a vital role in citrus canker development and pathogen spread ^[1, 3-5]^. The inhibition or disruption of pustule development can efficiently repressed pathogen spread and even confer plant resistance to citrus canker ^[6, 7]^, indicating that the manipulation of pustule development is a potential strategy for the efficient management of citrus canker. Thus, understanding the molecular mechanisms involved in responding to pathogen-induced pustule formation in citrus could stimulate renewed efforts to develop more effective and economical control methods of citrus canker management.

Auxin, a critical plant hormone that controls a range of plant growth and developmental processes, including cell division and expansion, has long been recognized as a regulator of plant defenses ^[8, 9]^. The effector AvrRpt2 from *Pseudomonas syringae* elicits auxin biosynthesis in plants and promotes disease in *Arabidopsis thaliana* ^[10]^. The flagellin Flg22 from *P. syringae* induces the microRNA mi393 to degrade the RNAs of the auxin receptor gene *TIR1*, resulting in immune responses in *Arabidopsis* ^[11]^. Auxin represses the expression of pathogenesis-related (*PR*) genes to impair defense responses, and the inhibition of auxin signaling is a part of the salicylic acid (SA)-mediated resistance mechanism ^[12-14]^. Indole-3-acetic acid (IAA) is the major form of auxin in most plants, including citrus. Many plant pathogens can either secret IAA into host tissues or enhance the host’s IAA synthesis, and the elevated IAA levels increase cell wall loosening and remodeling to favor pathogen invasion and spread ^[9]^. Conversely, plant disease resistance is enhanced by repressing auxin signaling or decreasing the IAA content ^[5, 15]^.

Xcc increases cell division and expansion in host-infected sites through the regulation of auxin to increase bacterial growth ^[3, 5]^. In the initial stages of canker development, naphthalene acetic acid treatments significantly enhanced the water soaking phenomenon on citrus leaves ^[5]^. Additionally, Costacurta et al. ^[16]^ reported that the Xcc pathogen produces IAA through the indole-3-pyruvic acid pathway, and that this IAA biosynthesis is enhanced by citrus leaf extracts. However, the molecular mechanisms of the host governing auxin responses to citrus canker remain to be elucidated.

The normal physiologic function of auxin depends on the spatiotemporal fine-tuning of hormone levels. The maintenance of IAA homeostasis is regulated by several groups of auxin-responsive genes, including those of the auxin/IAA (*Aux/IAA*), Gretchen Hagen3 (*GH3*) and small auxin-up RNA (*SAUR*) families ^[17]^. GH3 is an amido synthetase that conjugates IAA to amino acids (such as Asp, Ala, and Phe), thus inactivating free IAA ^[18]^. GH3 proteins are classified into three groups in *Arabidopsis*. Group I has jasmonic acid (JA)–amino synthetase activity, whereas group II is able to catalyze IAA conjugation to amino acids ^[18]^. No adenylation activity on the substrates tested has been found for group III members. Some GH3 proteins conjugate SA to amino acids ^[18]^. At present, many *GH3* genes have been identified in bean, apple, maize, tomato, rice and *Medicago sativa* ^[19-21]^. In addition to their functions in plant growth and development, *GH3* genes participate in disease resistance. *AtGH3.12* regulates SA-dependent defense responses by controlling pathogen-inducible SA levels ^[22]^. *AtGH3.5* has a dual regulatory role in *Arabidopsis* SA and auxin signaling during pathogen infection ^[13, 23]^. *Oryza sativa GH3-8* and *GH3-1* promote fungal resistance through the regulation of auxin levels ^[24, 25]^, while *GH3-2* mediates a broad-spectrum resistance to bacterial and fungal diseases ^[15]^.

In the early stage of this experiment, the transcriptomes of Newhall navel orange (*C. sinensis* Osbeck) and Calamondin (*Citrus madurensis*), with susceptibility and resistance to Xcc, respectively, were constructed (Unpublished). The transcriptomic analysis showed that GH3 group II *CsGH3.1* genes were induced significantly by Xcc and had high expression levels in the Newhall navel orange ^[26]^, indicating that this group’s members play important roles in responding to citrus canker. Here, to understand the roles of *CsGH3.1* in regulating host responses to citrus canker, we constructed transgenic Wanjingcheng orange (*C. sinensis* Osbeck) plants independently overexpressing *CsGH3.1* and *CsGH3.1L*. Their overexpression reduced endogenous auxin levels, altered plant architecture and enhanced host defense responses to citrus canker. We then explored the effects of *CsGH3.1* and *CsGH3.1L* overexpression in transgenic plants using high-throughput transcriptome sequencing.

## Materials and methods

### Plant materials and growth conditions

Wanjincheng orange (*C. sinensis* Osbeck) used in this study were planted in a greenhouse at the National Citrus Germplasm Repository, Chongqing, China.

### Vector construction

The coding sequences of the *CsGH3.1* (Cs1g22140) and *CsGH3.1L* (Cs8g04610) genes were obtained from the *C. sinensis* genome database (http://citrus.hzau.edu.cn/orange/). The pGEM plasmids independently containing the *CsGH3.1* and *CsGH3.1L* genes ^[26]^ and the plant expression vector pGN ^[27]^, from our laboratory, were used to construct plant overexpression vectors for this study. *CsGH3.1* and *CsGH3.1L* were digested from the pGEM vectors with *Sal*I/*Bam*HI, and then independently inserted into *Sal*I/*Bam*HI-digested pGN. Finally, two plant overexpression vectors containing *CsGH3.1* and *CsGH3.1L*, respectively, were obtained (Fig. S1). They were transformed into *Agrobacterium tumefaciens* strain EHA105 by electroporation.

### Citrus transformation

The epicotyls of Wanjincheng orange were used as explants for citrus transformation. The transformation protocol was performed according to Peng et al. ^[6]^. Putative transgenic shoots were screened using GUS histochemical staining ^[27]^. The recovery of GUS-positive plants was performed by grafting onto Troyer citrange [*Poncirus trifoliata* (L.) Raf. × *C. sinensis* Osbeck] seedlings *in vitro*. The integration of *CsGH3.1* and *CsGH3.1L* into plants was further confirmed by a PCR analysis. The primers for the PCR confirmation of transgenic plants are presented in Table S1. All transgenic and wild-type (WT) plants were grown in a netted greenhouse at 28°C.

### Real-time quantitative PCR (qPCR) analysis

Citrus total RNA was isolated using the EASYspin Plant RNA Extraction Kit following the manufacturer’s instructions (Aidlab, Beijing, China). RNA was reverse transcribed into cDNA using the iScriptTM cDNA Synthesis Kit (Bio-Rad, Hercules, CA, USA). The detection of gene expression was performed by qPCR using the iQ™ SYBR Green Supermix (Bio-Rad). The PCR reactions were carried out as follows: a pretreatment (94°C for 5 min), followed by 40 amplification cycles (94°C for 20 s and 60°C for 60 s). Experiments were repeated three times. Using the citrus Actin gene for normalization, the relative expression levels were calculated by the 2^−ΔΔCt^ method [28].

### Measurement of hormone contents

Hormones IAA, SA, zeatin (ZT), abscisic acid (ABA), ethylene (ET) and JA were extracted from the leaves of citrus plants was determined at Chongqing Bono Heng Biotechnology Co., Ltd (Chongqing, China). Contents of IAA, SA, ZT, ABA, and JA was simultaneously measured as described by ^[29, 30]^. In brief, Tissue samples (1 g fresh weight) were frozen in liquid nitrogen, ground to a fine powder, extracted with 80% methanol overnight and then centrifuged at 13,000 ×g for 10 min. The supernatant was evaporated and then redissolved in 1% acetic acid. Hormones were purified on Oasis cartridges (Waters, Milford, MA, USA) in accordance with the manufacturer’s instructions. The extracted hormones were redissolved in 10% methanol, and then IAA, SA, ZT, ABA and JA levels were determined using high-performance liquid chromatography. To measure ET contents, 1 g leaf tissue were placed in a gas-tight container equipped with septa, and 1 mL of headspace gas was sampled using a gas syringe ^[31]^. The ethylene production was measured using gas chromatograph. The test was repeated three times.

### Evaluation of transgenic plants resistance to citrus canker

The disease resistance assay for transgenic plants against citrus canker was performed according to Peng et al. ^[6]^. A Xcc strain, XccYN1 ^[6]^, was used to investigate plant disease resistance. Three mature healthy leaves per plant were tested. In total, 24 punctures were made per leaf with a needle containing the bacterial suspension (0.5 × 10^5^ CFU ml^−1^). The inoculated leaves were maintained at 28°C under a 16-h light/8-h dark photoperiod with 45 μmol m^−2^ s^−1^ illumination and 60% relative humidity. Photographs were taken at 10 dpi. The area of all diseased spots was assessed with ImageJ 2.0 software (National Institutes of Health, Bethesda, MD, USA). The disease intensity of an individual line was based on the mean values of the diseased areas surrounding the 24 punctures on three leaves using the rating index described by Peng et al. ^[6]^. The test was repeated three times.

### Construction of RNA-Seq libraries and high-throughput sequencing

In this experiment, three biological replicates per selected transgenic and WT plants were performed. Total RNA from fully mature leaves was extracted using a TRIzol kit (Invitrogen, Thermo Fisher Scientific, Shanghai, China) in accordance with the user manual. RNA quality was determined using a NanoDrop 2000 spectrophotometer (Thermo) and an Agilent 2100 Bioanalyzer (Agilent Technologies Canada Inc, Mississauga, ON, Canada). Sequencing libraries were constructed from 1 µg of total RNA using NEBNext Ultra^™^ RNA Library Prep Kit for Illumina (New England Biolabs, Ipswich, MA, USA) following manufacturer’s recommendations. The libraries were sequenced using the Illumina HiSeq 2500 platform (BioMarker Technologies Illumina, Inc, Shanghai, China).

### Analysis of RNA-Seq data

Approximately 6.4 Gb of high-quality clean reads were generated from each library after removing adaptor sequences and filtering low-quality sequences. All the clean reads were mapped to the reference genome of sweet orange (*C. sinensis*, http://citrus.hzau.edu.cn/orange/index.php) by TopHat2 with default parameters ^[32]^. Gene function was annotated based on Nr, Nt, Pfam, KOG/COG, Swiss-Prot, KO and gene ontology (GO) databases. Gene expression levels in all the biological replicates were estimated using the FPKM method ^[33]^. A differential expression analysis between transgenic lines and WT was performed using the DESeq R package 1.10.1^[34]^. Genes with adjusted P-values < 0.05 found by DESeq were assigned as differentially expressed genes (DEGs). A GO enrichment analysis of the DEGs was performed using the GOseq R package ^[35]^. A KEGG pathway enrichment analysis of DEGs was performed using KOBAS software ^[36]^.

MapMan software ^[37]^ was also used to analyze citrus gene expression data. At the end, the citrus genes from the reference genome of sweet orange (*C. sinensis*, http://citrus.hzau.edu.cn/orange/index.php) were assigned to BINs using the Mercator automated annotation pipeline (http://mapman.gabipd.org/web/guest/mercator), and then, the pathways, which were affected by DEGs, were analyzed using MapMan.

Differentially represented MapMan pathways were defined using a two-tailed Wilcoxon rank sum test corrected using the Benjamin–Hochberg method (false discovery rate ⩽ 0.05).

## Results

### Overexpression of *CsGH3.1* and *CsGH3.1L* in citrus causes abnormal phenotypes

To understand the functions of *CsGH3.1* and *CsGH3.1L* in citrus responses to citrus canker, the two genes, under the control of CaMV 35S promoter (Fig. S1), were introduced separately into Wanjincheng orange by *Agrobacterium*-mediated epicotyl transformation. Transgenic plants were identified using β-glucuronidase (GUS) histochemical staining and PCR analysis (Fig. S1). Expression levels of *CsGH3.1* and *CsGH3.1L* in transgenic plants were evaluated by qPCR analyses (Fig. S2). Based on the data, the pLGN-GH3.1 lines 1-3, -4, -5, -6, -8 and -9, and the pLGN-GH3.1L lines L-2, -5 and -6 showed high expression levels.

After the transgenic lines were planted in a greenhouse, their phenotypes were investigated. In the early stage (∼6 months after planting), most transgenic plants showed leaf drooping and upward curling (Fig. 1a). Gradual increased branching and leaf curling were detected as transgenic plants grew (Fig. S3). Lines 1-3, 1-4, 1-5, 1-9, L-2, L-5 and L-6, having high gene expression levels, had severe malformations, while lines 1-1, 1-2 and 1-10, having low gene expression levels, showed no obvious differences compared with WT (Fig. S3). Line 1-8 also showed no obvious difference, while line L-1 displayed mild changes in its phenotype compared with WT (Fig. S3).

**Fig. 1.**
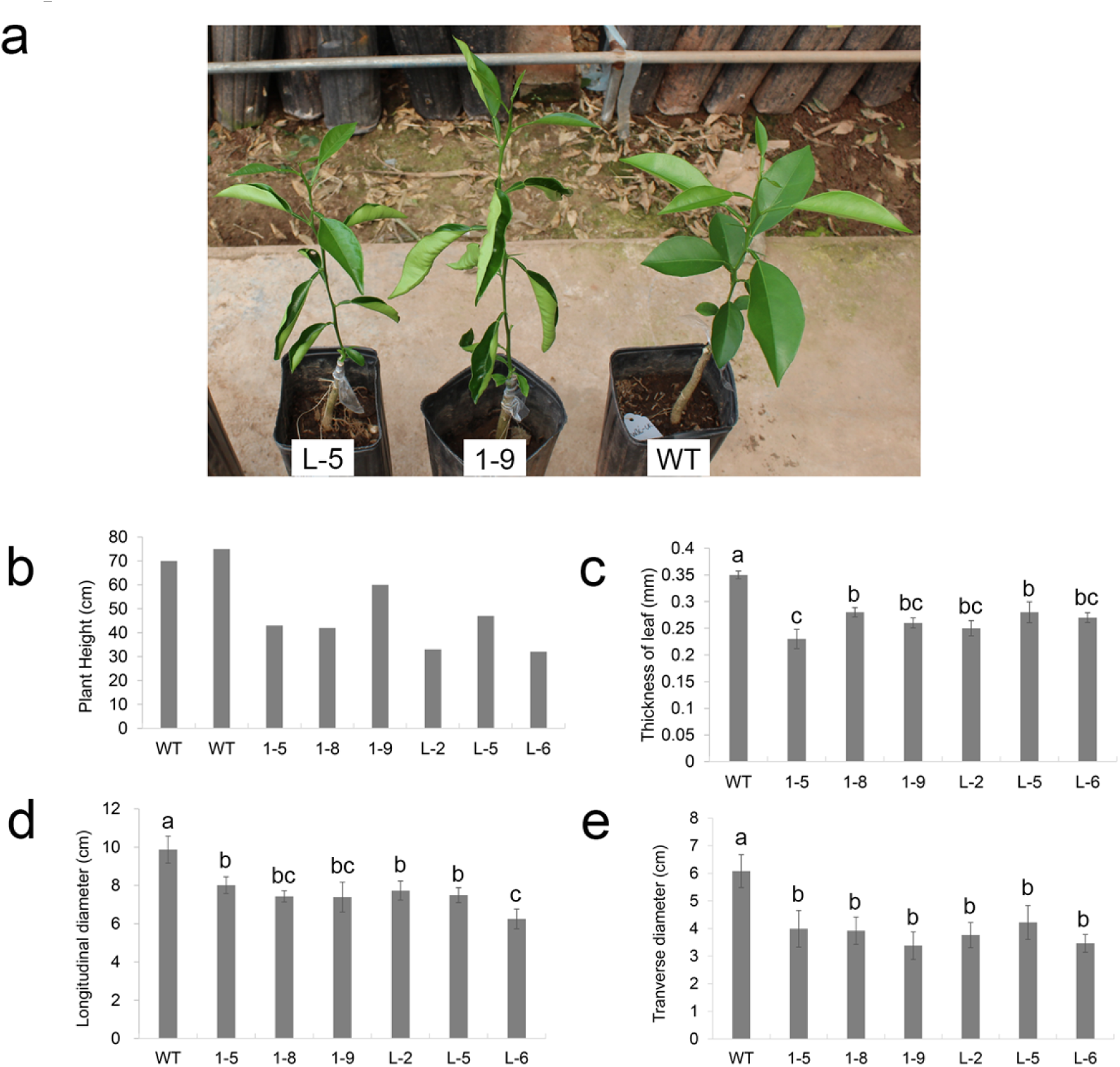
Phenotypic analysis of transgenic citrus independently overexpressing *CsGH3.1* (1-#) and *CsGH3.1L* (L-#). (**a**) Phenotypes of transgenic and wild-type (WT) plants after growing for six months in a greenhouse. (**b**) The heights of transgenic plants grown in a greenhouse for two years after grafting. The thickness (**c**), and the longitudinal (**d**) and transverse (**e**) diameters of leaves from transgenic plants grown in a greenhouse for 2 years after grafting were evaluated using 20 leaves per line. Error bars represent the mean standard errors. Different letters on top of the bars represent significant differences from WT controls based on a Tukey’s test (*P* < 0.05).

After one year, lines 1-3 and 1-4 died. After two years, transgenic plants displayed a bushy dwarf phenotype with smaller, drooping and upward curling leaves and branch softening and drooping (Fig. 1b and Fig. S4). Moreover, transgenic leaves were significantly thinner, and both their longitudinal and transverse diameters were significantly shorter compared with WT (Fig. 1c–e). Such abnormal phenotypes indicated that overexpressing *CsGH3.1* and *CsGH3.1L* affected the basic plant development.

### The decreased free IAA level in transgenic plants

To investigate the auxin content, the free IAA level in each transgenic line was measured. The free IAA levels in the 1-5, 1-8, 1-9, L-2, L-5 and L-6 transgenic lines were significantly lower than in WT plants before exposure to Xcc (Fig. 2). Other lines showed no differences in free IAA levels compared with WT. After Xcc inoculation, free IAA levels in these transgenic lines were still significantly lower than in WT, although the levels were markedly decreased in both transgenic lines and WT after Xcc inoculations (Fig. 2).

**Fig. 2.**
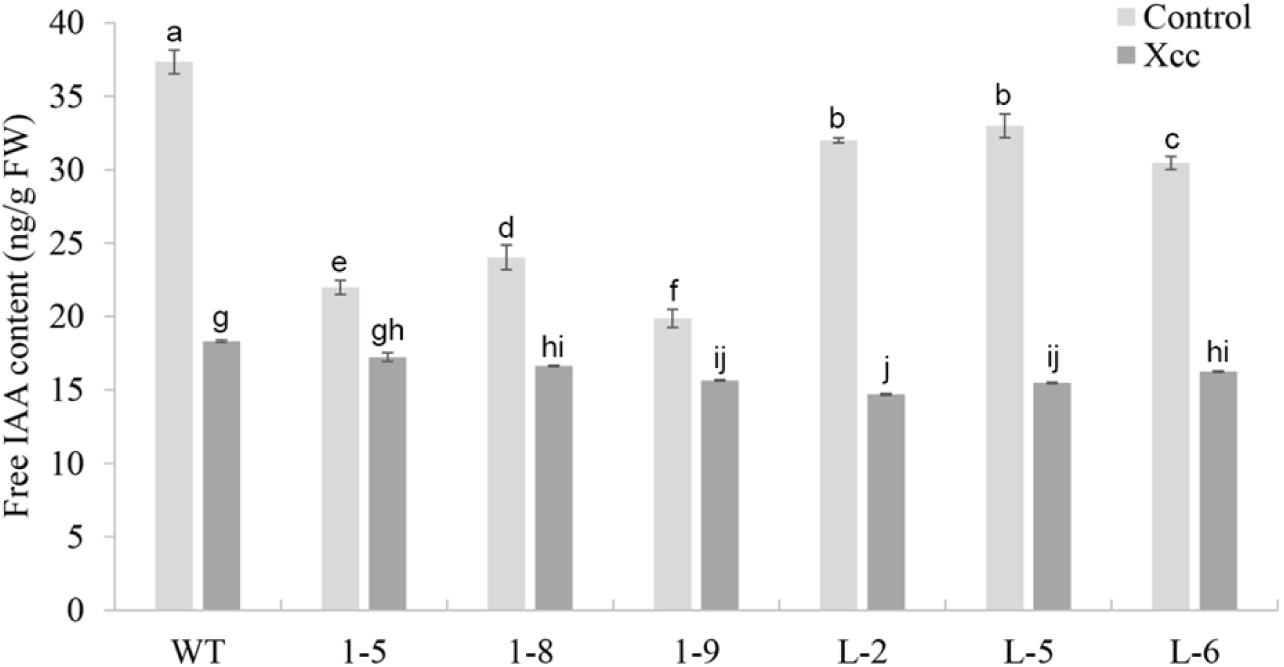
Determination of the IAA contents in transgenic citrus independently overexpressing *CsGH3.1* (1-#) and *CsGH3.1L* (L-#). IAA was isolated from six fully expanded intact leaves per line. The presented IAA values are the averages of three independent measurements per line before *Xanthomonas citri* subsp. *citri* (Xcc) infection (control) and 3 d after Xcc infection. The overexpression of *CsGH3.1* or *CsGH3.1L* decreased the IAA contents in transgenic plant before and after pathogen exposure. Error bars represent the mean standard errors. WT: wild-type. Different letters on top of bars represent significant differences from WT controls based on Tukey’s test (*P* < 0.05).

### Overexpression of *CsGH3* in citrus reduced susceptibility to Xcc

To evaluate the citrus canker resistance levels of transgenic plants, the 1-5, 1-8, 1-9, L-2, L-5 and L-6 transgenic lines were inoculated with Xcc by *in vitro* pinpricks. The diseased areas were determined 10 d after Xcc infection. Lesions around the pinprick sites in transgenic lines were significantly smaller compared with WT (Fig. 3a and b). The disease indices of these transgenic lines decreased significantly compared with WT (Fig. 3c), indicating that these transgenic lines had enhanced citrus canker resistance. The lines 1-9 and L-5 showed the strongest resistance to citrus canker (Fig. 3c). Thus, overexpressing *CsGH3.1* and *CsGH3.1L* enhanced host defense responses to citrus canker.

**Fig. 3.**
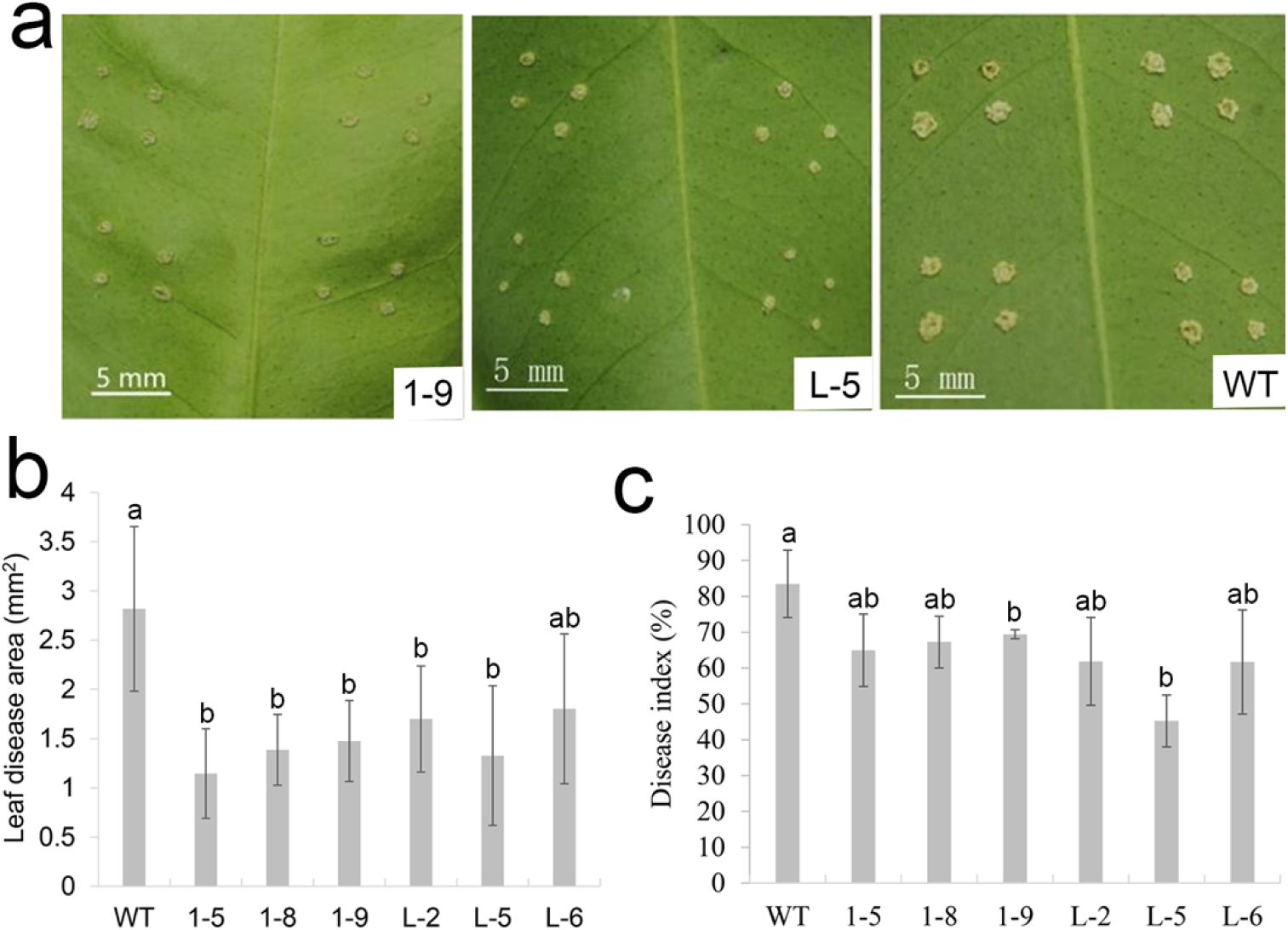
Evaluation of citrus canker resistance in transgenic citrus independently overexpressing *CsGH3.1* (1-#) and *CsGH3.1L* (L-#). (**a**) Citrus canker leaf symptoms in transgenic and non-transgenic lines 10 d after pin-puncture inoculation with *Xanthomonas citri* subsp. *citri* (Xcc). (**b**) Lesion areas and (**c**) and disease indices in transgenic plants. Diseased areas in leaves were counted 10 d after Xcc inoculation in WT and transgenic lines. Each column represents the mean of nine leaves from three independent experiments. Error bars represent the mean standard errors. Different letters on top of bars represent significant differences from wild-type (WT) controls based on Tukey’s test (*P* < 0.05).

### An overview of the transcriptional responses in transgenic citrus lines

To reveal the molecular mechanisms underlying canker resistance in CsGH3-overexpressing plants, global transcriptional profiling of lines 1-9 and L-5 showing high levels of resistance to citrus canker and a WT plant were compared using RNA-Seq (Fig. 4a; Supplementary data S1). In total, 1,560 and 1,037 genes were identified as DEGs in the 1-9 and L-5 transgenic lines, respectively, when compared with a WT plant (Supplementary data S2). There were more upregulated DEGs than downregulated DEGs in both the 1-9 and L-5 lines (Fig. 4b; Supplementary data S2). In the GO annotation, most of the DEGs were classified into metabolic process, cellular process, single-organism process, response to stimulus, and biological regulation (Supplementary data S3; Fig. S5). The KEGG pathway enrichment analysis further revealed that the DEGs in the 1-9 and L-5 transgenic lines were assigned to 88 and 83 KEGG pathways (Fig. 4c), respectively. Notably, 34 and 31 DEGs were significantly (q-value < 0.05) assigned to “plant hormone signaling transduction (KO04075)” (Fig. 4c).

**Fig. 4.**
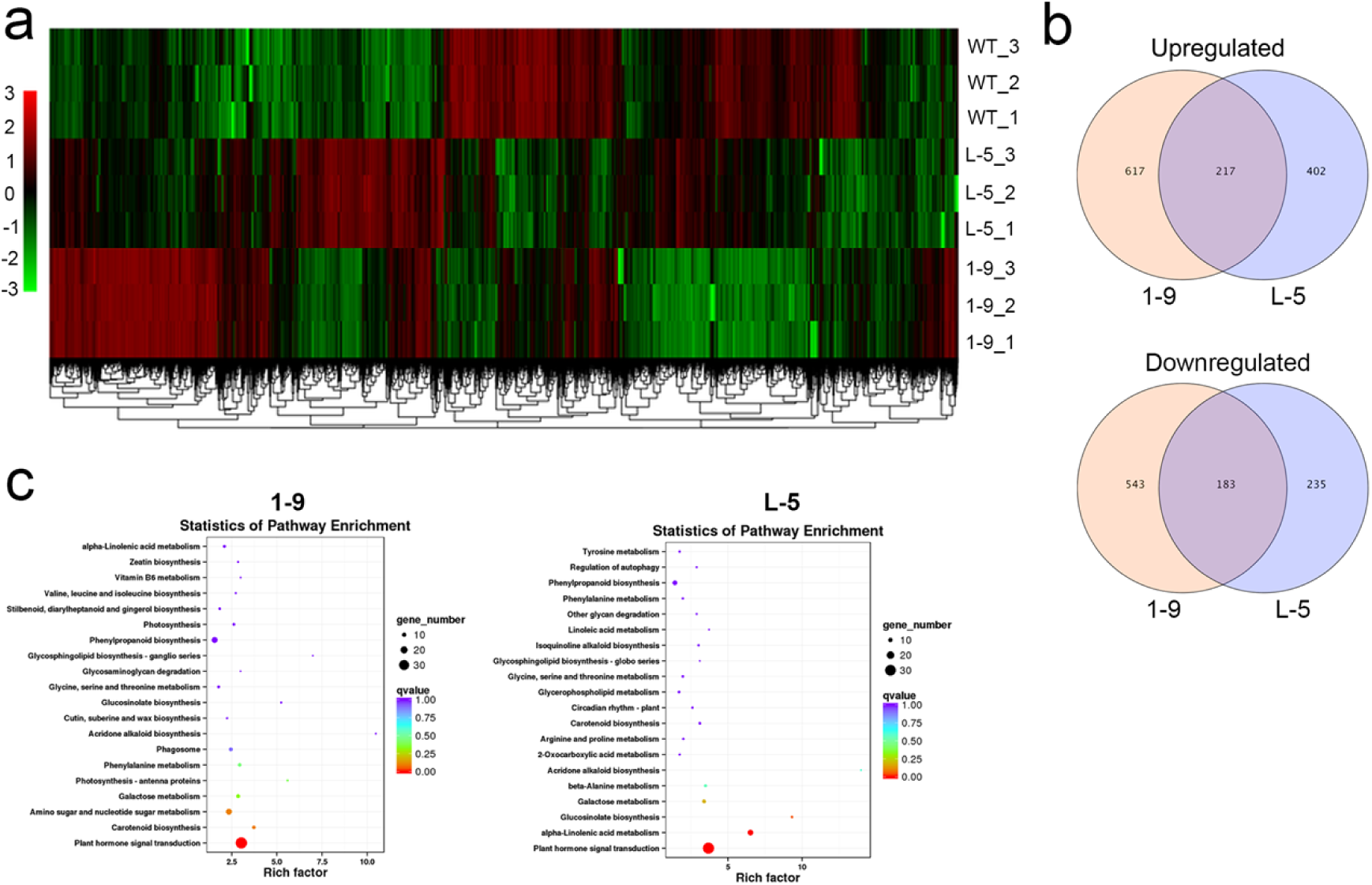
Global gene expression profiles of transgenic citrus independently overexpressing *CsGH3.1* (1-9) and *CsGH3.1L* (L-5). (**a**) Heat map analysis of the differentially expressed genes (DEGs) among 1-9 and L-5 transgenic and wild-type lines. The 1-9 and L-5 transgenic lines showed a similar hierarchical cluster pattern. (**b**) Venn diagrams showing the overlaps of differentially expressed genes between transgenic lines. In total, 400 of 2,745 DEGs showed similar expression profiles among these lines. (**c**) KEGG pathway enrichment of the DEGs between transgenic lines. The overexpression of Cs*GH3.1* or Cs*GH3.1L* significantly affected hormone signal transduction.

To further survey the pathways or functions that were affected by the DEGs in the 1-9 and L-5 transgenic lines, the RNA-Seq data were visualized using the MapMan tool (Fig. 5). A complete list of MapMan pathways differentially represented in the transgenic lines is provided in Supplementary data S4. Based on these data, cell wall, secondary metabolism, hormone metabolism, stress, RNA and signaling were the major pathways or functions that were significantly regulated by the overexpression of *CsGH3.1* and *CsGH3.1L*. Differences in differentially represented pathways or functions were also detected between the transgenic lines. Importantly, flavonol metabolism, cytokinin metabolism and synthesis-degradation, biotic stress, touch or wounding, Aux/IAA family, signaling, and unknown categories displayed significant changes in both the 1-9 and L-5 transgenic lines compared with WT. Biotic stress-related pathways were positively affected, while Aux/IAA family genes were negatively affected by both CsGH3.1 and CsGH3.1L (Fig. 5). Aux/IAA family genes have vital roles in auxin signaling ^[9]^.The survey clearly showed that the overexpression of *CsGH3.1* and *CsGH3.1L* repressed auxin signaling and enhanced biotic stress responses in citrus. Thus, the DEGs involved in auxin metabolism and signaling, as well as biotic stress responses, were studied in more detail.

**Fig. 5.**
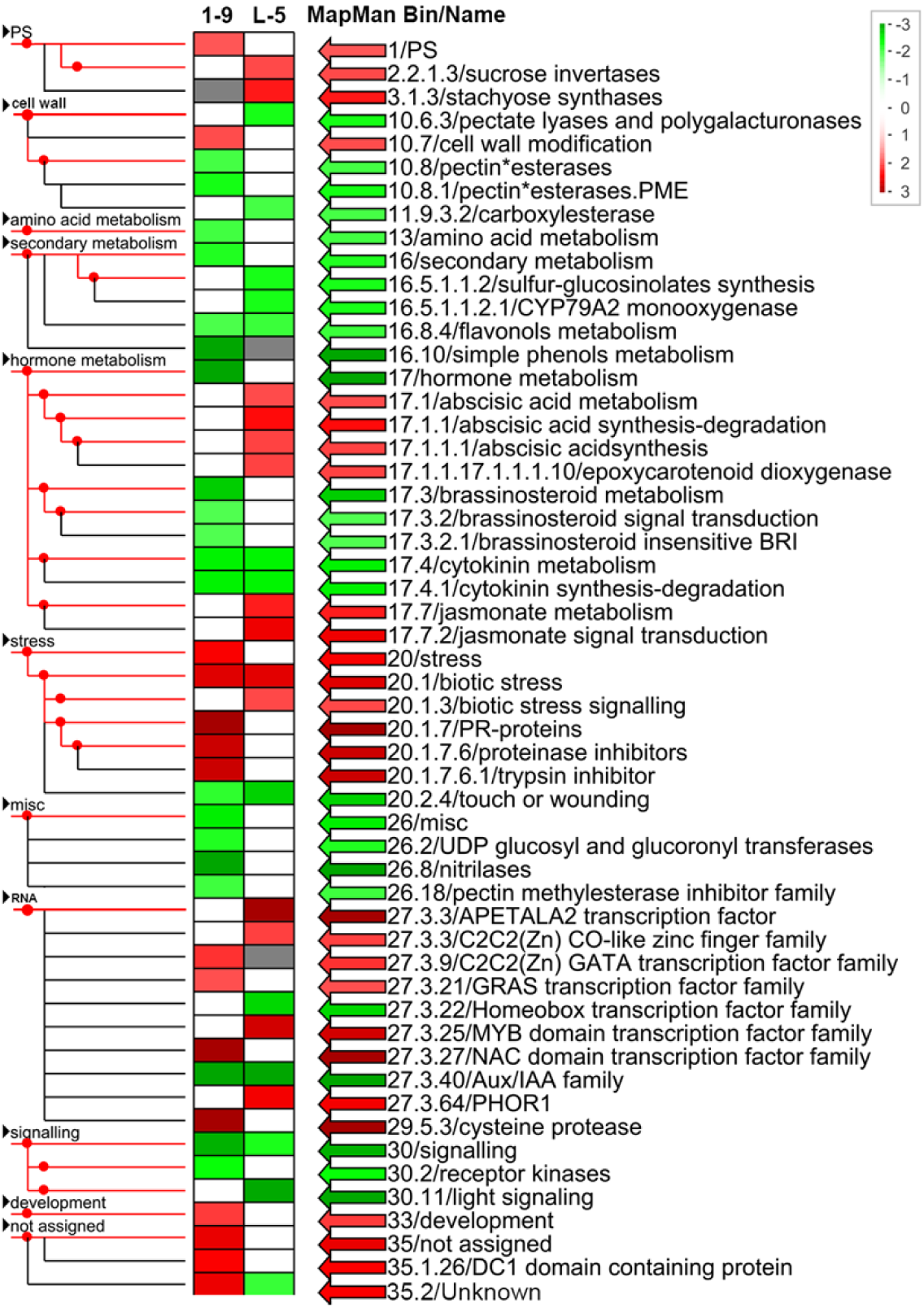
MapMan visualization of differentially represented pathways and functional categories between the 1-9 and L-5 transgenic citrus lines overexpressing *CsGH3.1* and *CsGH3.1L*, respectively. Each colored rectangular block denotes a MapMan pathway or functional category. Upregulated and downregulated categories are shown in red and green, respectively. The scale bar represents fold change values. The categories differentially represented in the transgenic plants are indicated on the right.

### Auxin-related genes

We investigated the DEGs related to auxin homeostasis, perception and signaling in transgenic lines using a MapMan analysis. Most of the 28 auxin-related DEGs showed significantly downregulated expression levels in both the 1-9 and L-5 transgenic lines, and most of these genes were assigned to auxin signaling transduction (Table 1). All 12 Aux/IAA family members, a group of auxin-induced genes, showed significantly downregulated expression levels in the transgenic plants.

**Table 1.**
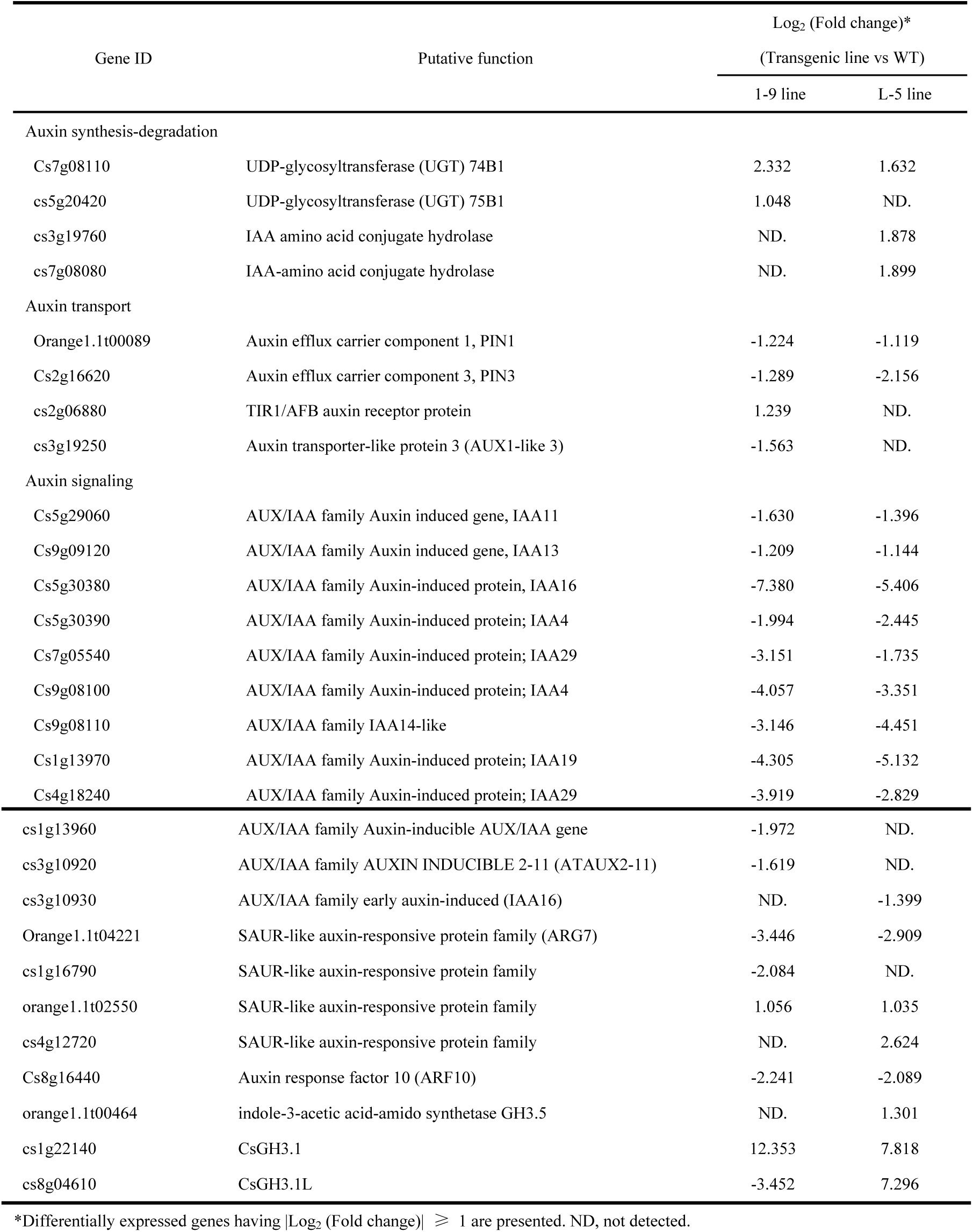
Differentially expressed genes related to auxin biosynthesis and signaling in transgenic citrus independently overexpressing *CsGH3.1* and *CsGH3.1*

| Gene ID | Putative function | Log<sub>2</sub> (Fold change)*<br>(Transgenic line vs WT) |  |
| --- | --- | --- | --- |
|  |  | 1-9 line | L-5 line |
| Auxin synthesis-degradation |  |  |  |
| Cs7g08110 | UDP-glycosyltransferase (UGT) 74B1 | 2.332 | 1.632 |
| cs5g20420 | UDP-glycosyltransferase (UGT) 75B1 | 1.048 | ND. |
| cs3g19760 | IAA amino acid conjugate hydrolase | ND. | 1.878 |
| cs7g08080 | IAA-amino acid conjugate hydrolase | ND. | 1.899 |
| Auxin transport |  |  |  |
| Orange1.1t00089 | Auxin efflux carrier component 1, PIN1 | -1.224 | -1.119 |
| Cs2g16620 | Auxin efflux carrier component 3, PIN3 | -1.289 | -2.156 |
| cs2g06880 | TIR1/AFB auxin receptor protein | 1.239 | ND. |
| cs3g19250 | Auxin transporter-like protein 3 (AUX1-like 3) | -1.563 | ND. |
| Auxin signaling |  |  |  |
| Cs5g29060 | AUX/IAA family Auxin induced gene, IAA11 | -1.630 | -1.396 |
| Cs9g09120 | AUX/IAA family Auxin induced gene, IAA13 | -1.209 | -1.144 |
| Cs5g30380 | AUX/IAA family Auxin-induced protein, IAA16 | -7.380 | -5.406 |
| Cs5g30390 | AUX/IAA family Auxin-induced protein; IAA4 | -1.994 | -2.445 |
| Cs7g05540 | AUX/IAA family Auxin-induced protein; IAA29 | -3.151 | -1.735 |
| Cs9g08100 | AUX/IAA family Auxin-induced protein; IAA4 | -4.057 | -3.351 |
| Cs9g08110 | AUX/IAA family IAA14-like | -3.146 | -4.451 |
| Cs1g13970 | AUX/IAA family Auxin-induced protein; IAA19 | -4.305 | -5.132 |
| Cs4g18240 | AUX/IAA family Auxin-induced protein; IAA29 | -3.919 | -2.829 |
| cs1g13960 | AUX/IAA family Auxin-inducible AUX/IAA gene | -1.972 | ND. |
| cs3g10920 | AUX/IAA family AUXIN INDUCIBLE 2-11 (ATAUX2-11) | -1.619 | ND. |
| cs3g10930 | AUX/IAA family early auxin-induced (IAA16) | ND. | -1.399 |
| Orange1.1t04221 | SAUR-like auxin-responsive protein family (ARG7) | -3.446 | -2.909 |
| cs1g16790 | SAUR-like auxin-responsive protein family | -2.084 | ND. |
| orange1.1t02550 | SAUR-like auxin-responsive protein family | 1.056 | 1.035 |
| cs4g12720 | SAUR-like auxin-responsive protein family | ND. | 2.624 |
| Cs8g16440 | Auxin response factor 10 (ARF10) | -2.241 | -2.089 |
| orange1.1t00464 | indole-3-acetic acid-amido synthetase GH3.5 | ND. | 1.301 |
| cs1g22140 | CsGH3.1 | 12.353 | 7.818 |
| cs8g04610 | CsGH3.1L | -3.452 | 7.296 |

\*Differentially expressed genes having $|\text{Log}_2(\text{Fold change})| \geq 1$ are presented. ND, not detected.

In addition, two of four SAUR-like auxin-responsive protein family members (ARG7), and one auxin response factor (ARF10) also showed significantly downregulated expression levels in the transgenic plants. The expression levels of three genes (PIN1, PIN3 and AUX1-like protein 3) involved in auxin transport were significantly downregulated in transgenic plants (Table 1). However, four genes (Cs7g08110, Cs5g20420, Cs3g19760 and Cs7g08080) involved in auxin synthesis-degradation were significantly induced by the independent overexpression of *CsGH3.1* and *CsGH3.1L* (Table 1). Overall, the overexpression of *CsGH3.1* and *CsGH3.1L* significantly repressed the expression levels of auxin transport and signaling-related genes. Interestingly, *CsGH3.1*’s overexpression downregulated *CsGH3.1L*’s expression, while *CsGH3.1L*’s overexpression upregulated *CsGH3.1*’s expression (Table 1).

### Disease response-related genes

Figure 6 presents an overview of the MapMan functional categories for genes involved in disease responses. Their functional products could be sorted into the following major classes: hormone signaling, cell wall, proteolysis, signaling, PR proteins, redox state, transcription factors, heat shock proteins and secondary metabolites. Among the disease response-related genes, the number (90) of upregulated genes was obvious more than that (57) of downregulated genes. In total, 45 and 23 genes, including resistance, stress recognition, signal receptor and transduction, and PR protein, were directly assigned into the “stress. biotic” MapMan category in the 1-9 and L-5 transgenic lines, respectively (Supplementary data S5). Among these genes, 33 and 18 genes showed significantly upregulated expression levels in the 1-9 and L-5 lines, respectively (Supplementary data S5).

**Fig. 6.**
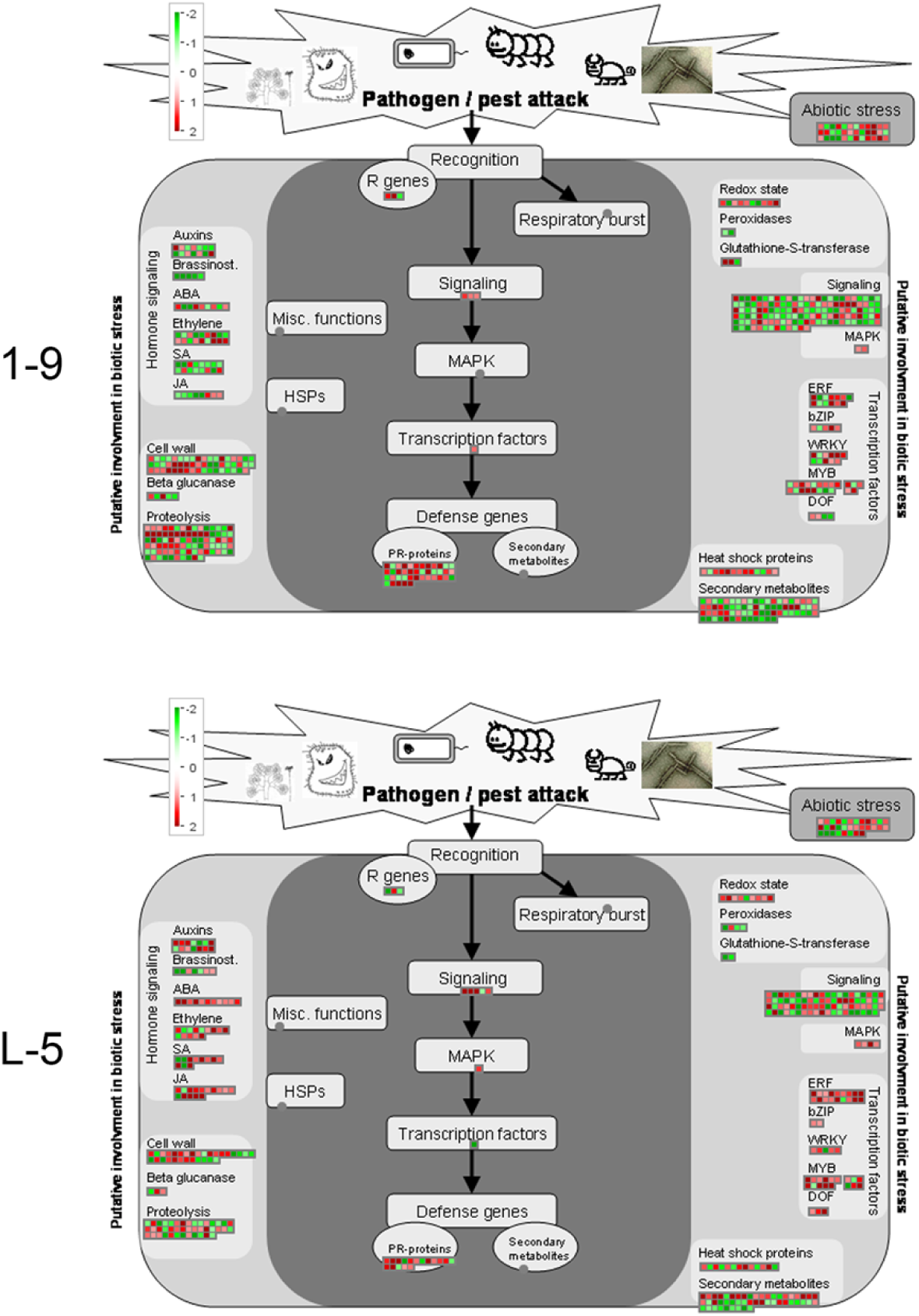
MapMan visualization of the functional categories of genes differentially expressed in response to biotic stress in the 1-9 and L-5 transgenic citrus lines overexpressing *CsGH3.1* and *CsGH3.1L*, respectively. Significantly upregulated and downregulated genes are displayed in red and green, respectively.

Table 2 displays the important DEGs correlated with biotic stress in transgenic plants based on the MapMan annotation. The induced genes included a Toll-Interleukin-Resistance (TIR) domain family member (Cs5g18230) involved in innate immune responses, three defense *PR* genes (Cs6g01070, NB-ARC domain-containing disease resistance protein; Cs9g18740, disease resistance family protein/LRR family protein; and Cs5g19240, TIR-NBS-LRR disease resistance protein). Moreover, all 10 transcription factor genes had upregulated expression levels, and these genes mainly included the AP2/EREPB, MYB and WRKY families. Thus, gene expression associated with responses to biotic stress was clearly activated by the overexpression of *CsGH3.1* and *CsGH3.1L*.

**Table 2.** Differentially expressed genes related to biotic stress in transgenic citrus independently overexpressing

*CsGH3.1* and *CsGH3.1L*
| Gene ID | Putative function | Log<sub>2</sub> Fold change*<br>(Transgenic line vs WT) |  |
| --- | --- | --- | --- |
|  |  | 1-9 | L-5 |
| Stress recognition |  |  |  |
| Orange1.1t02076 | Disease resistance-responsive family protein | -1.005 | -2.029 |
| Orange1.1t03601 | LRR receptor-like kinase FLS2 | 2.755 | 1.412 |
| Cs2g16870 | MLP-like protein 31 (MLP31) | 2.198 | 3.223 |
| Cs2g17820 | ARM repeat superfamily protein | 1.177 | 3.669 |
| Cs2g29780 | HXXXD-type acyl-transferase family protein | 1.436 | 3.488 |
| Cs2g31450 | Acetyl coa:(Z)-3-hexen-1-ol acetyltransferase (CHAT) | -1.365 | 1.685 |
| Signal receptor |  |  |  |
| Cs5g18230 | Toll-Interleukin-Resistance (TIR) domain family protein | 2.044 | 1.717 |
| Cs5g27950 | Leucine-rich repeat I receptor kinase | -2.207 | -2.005 |
| Cs5g15420 | Leucine-rich repeat VIII (VIII-2) receptor kinase | 1.262 | 1.429 |
| Cs9g14980 | Leucine-rich repeat XI receptor kinase | -1.142 | -2.069 |
| Orange1.1t01442 | Leucine-rich repeat XI receptor kinase | -2.353 | -1.839 |
| Orange1.1t03106 | Leucine-rich repeat XI receptor kinase | 1.394 | 1.342 |
| Cs4g07400 | Serine/threonine receptor kinase | 3.036 | 1.785 |
| Orange1.1t01406 | DUF 26receptor kinase | -1.357 | -1.596 |
| Signal transduction |  |  |  |
| Cs1g17210 | JAZ1 involved in jasmonate signaling | 1.173 | 2.883 |
| Cs1g17220 | JAZ1 involved in jasmonate signaling | 1.143 | 2.371 |
| Cs6g17590 | Calmodulin-like protein | 1.321 | 1.482 |
| Cs7g27120 | Calmodulin binding protein-like | 1.230 | 1.591 |
| Cs5g07160 | Calmodulin like 37 (CML37) | 1.288 | 1.996 |
| Cs2g15970 | Rac-like GTP-binding protein RAC2 | -2.151 | -1.370 |
| Cs2g21150 | Mitogen-activated protein kinase kinase kinase 18 (MAPKKK18) | 1.286 | 3.530 |
| Orange1.1t01873 | Tobacco Rapid Alkalinization Factor (RALF) | -1.357 | -1.438 |
| Cs9g05280 | AHP1 Arabidopsis thaliana histidine phosphotransfer protein | -1.134 | -1.235 |
| Cs7g09390 | Phototropic-responsive NPH3 family protein | -1.633 | -1.696 |
| Cs9g07670 | Chlorophyll A-B binding family protein | -1.136 | -2.031 |
| Defense response gene |  |  |  |
| Cs6g01070 | NB-ARC domain-containing disease resistance protein | 2.089 | 1.506 |
| Cs9g18740 | Disease resistance family protein / LRR family protein defense | 2.640 | 2.459 |
| Cs8g14950 | Glucan endo-1,3-beta-glucosidase | -1.567 | 1.177 |
| Cs3g06300 | Receptor like protein 6 (RLP6) | -1.089 | 1.200 |
| Cs5g19240 | Disease resistance protein (TIR-NBS-LRR class) | 2.180 | 1.563 |
| Redox metabolism |  |  |  |
| Cs2g17910 | Cytochrome b561/ferric reductase transmembrane protein family | -1.646 | -1.768 |

| Gene ID | Putative function | Log<sub>2</sub> Fold change* |  |
| --- | --- | --- | --- |
|  |  | (Transgenic line vs WT) |  |
| Cs5g32580 | Thioredoxin superfamily protein | 1.316 | 1.028 |
| Cs2g16150 | GRX480, the glutaredoxin family that regulates protein redox state | 1.436 | 1.806 |
| Orange1.1t03455 | Probable glutathione S-transferase, gsts , Auxin-induced protein | -1.777 | -1.624 |
| Cs2g15310 | Peroxidase superfamily protein | -1.918 | -1.385 |
| Responsive transcription factor |  |  |  |
| Cs6g15360 | The DREB subfamily A-1 of ERF/AP2 transcription factor (CBF4) | 2.032 | 4.256 |
| Cs9g16810 | The DREB subfamily A-1 of ERF/AP2 transcription factor (CBF4) | 1.725 | 2.039 |
| Orange1.1t01154 | The DREB subfamily A-1 of ERF/AP2 transcription factor (CBF2) | 1.543 | 2.840 |
| Cs1g07950 | The ERF subfamily B-1 of ERF/AP2 transcription factor (ATERF-4) | 1.913 | 2.078 |
| Cs4g07040 | The DREB subfamily A-5 of ERF/AP2 transcription factor family | 5.413 | 2.803 |
| Cs5g29830 | Member of the R2R3 factor gene family (MYB14) | 1.249 | 2.333 |
| Cs2g27410 | Member of the R2R3 factor gene family (MYB58) | 3.896 | 4.470 |
| Cs3g23070 | Member of the R2R3 factor MYB gene family (MYBR1) | 2.909 | 2.012 |
| Cs3g23950 | Member of the R2R3 factor gene family (MYB73) | 2.235 | 3.114 |
| Cs1g03870 | Group II-c WRKY Transcription Factor (WRKY51) | 3.168 | 1.313 |
| Protein degradation |  |  |  |
| Cs2g18910 | Encode a protein similar to subtilisin-like serine protease | -1.763 | -2.831 |
| Cs2g27790 | Cysteine proteinases superfamily protein | 3.370 | 1.277 |
| Cs9g06150 | Eukaryotic aspartyl protease family protein | 1.971 | 3.298 |
| Cs2g29430 | Skp2-like F-box family protein | -2.364 | -2.155 |
| Cs3g14660 | F-box family protein; RNI-like superfamily protein | 1.338 | 1.078 |
| Cs6g13640 | E3 ubiquitin ligase protein involved in PAMP-triggered immunity | 1.070 | 2.153 |

\*Differentially expressed genes having |Log<sub>2</sub> (Fold change)| ≥ 1 are presented.

Data presented in Table 3 show that 11 cell wall-related genes were differentially expressed, with 8 being upregulated by the overexpression of *CsGH3*. These upregulated genes were mostly involved in cell wall biosynthesis and modification, such as xyloglucan endotransglycosylase, cellulose synthase, and wax and cutin synthesis. Among them, two genes encoding FASCICLIN-like arabinogalactan-protein 12 (Cs8g16830) and a xylem-specific cellulose synthase (Cs4g02000), showed highly induced expression levels (fold change > 5). Three genes (pectinesterase inhibitor Cs1g05510, expansin Cs5g07854 and polygalacturonase Cs7g08620), assigned to cell wall degradation, were downregulated when CsGH3 was overexpressed.

**Table 3.**
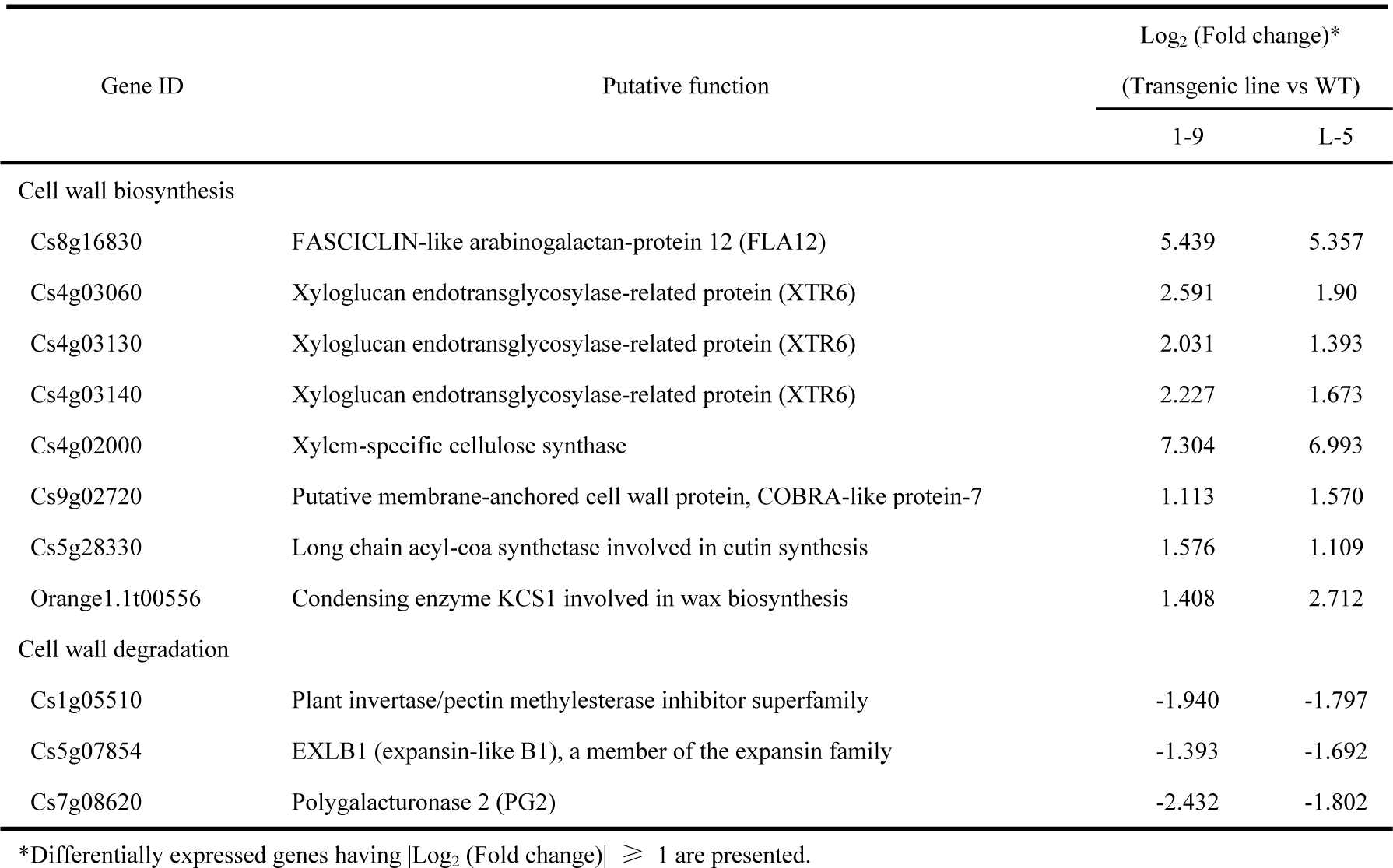
Differentially expressed genes related to cell wall in transgenic citrus independently overexpressing

### Effects of overexpression of *CsGH3* on hormone contents of transgenic lines

Because the transgenic plants displayed severe dwarfism (Fig. S4) and obvious changes in transcriptional profiles involved in hormone metabolism (Fig. 6; Supplementary data S6), we investigated SA, JA, ABA, ZT and ET levels in the 1-9 and L-5 transgenic lines (Fig. 7). The SA and ET contents in both 1-9 and L-5 transgenic lines were significantly greater than in WT. The JA, ABA and ZT contents in the 1-9 transgenic line were significantly lower than in WT. No significant difference in JA, ABA or ZT level was detected in the L-5 line compared with WT, although the ABA and ZT levels were decreased in this transgenic line.

**Fig. 7.**
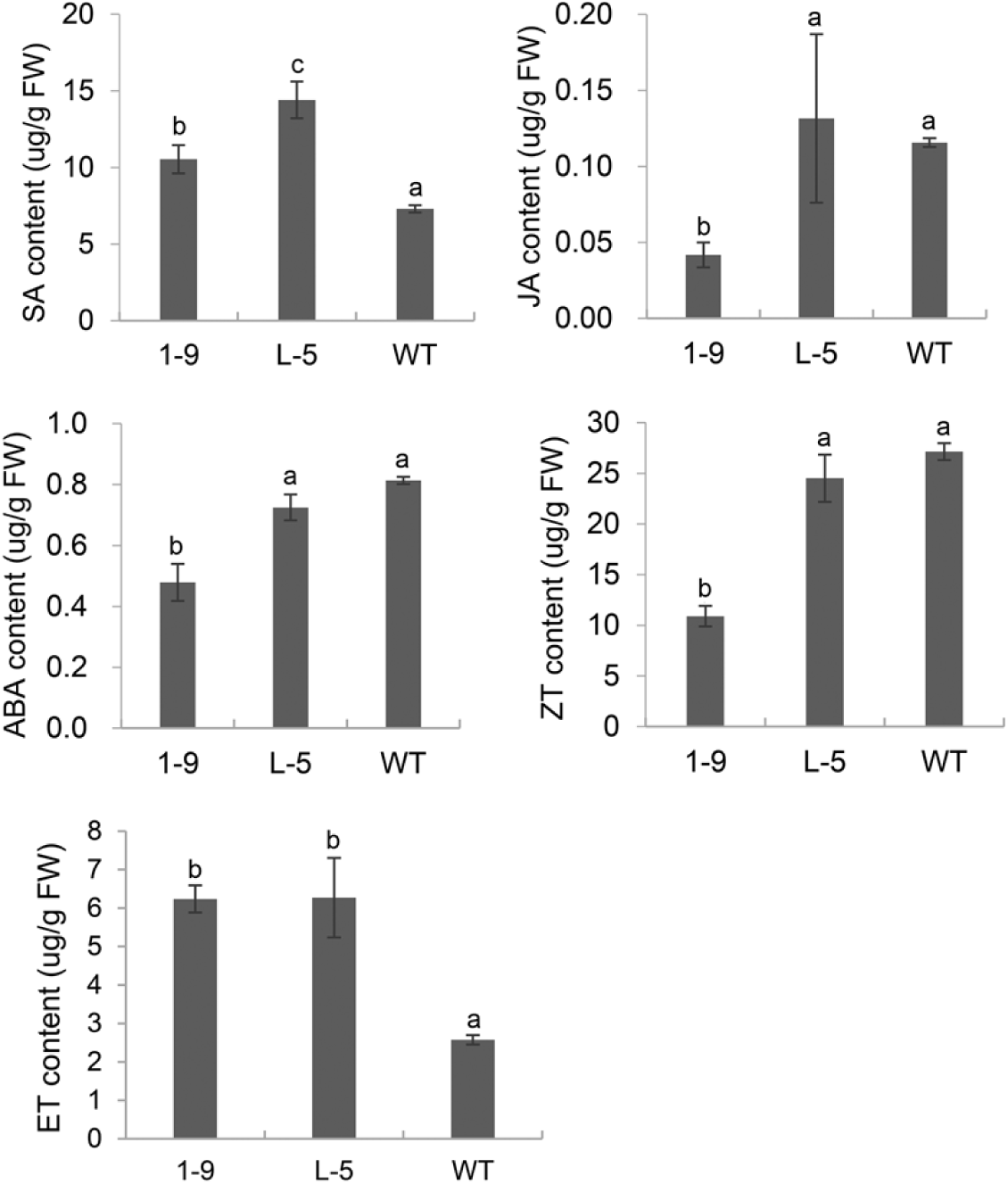
Determination of SA, JA, ABA, ZT and ET contents in the 1-9 and L-5 transgenic citrus lines overexpressing *CsGH3.1* and *CsGH3.1L*, respectively. Hormones were isolated from six fully expanded intact leaves per line. Error bars represent the mean standard errors of three independent measurements. Different letters on top of bars represent significant differences from wild-type (WT) controls based on Tukey’s test (*P* < 0.05).

## Discussion

The *GH3* gene family maintains hormonal homeostasis by conjugating free hormones to amino acids during exposure to biotic and abiotic stresses. Our work first showed that the overexpression of *CsGH3.1* and *CsGH3.1L* decreased free IAA levels. Correspondingly, transgenic plants displayed a bushy dwarf phenotype that is associated with an auxin shortage ^[25]^. Additionally, naphthalene acetic acid treatments significantly induced the expression of *CsGH3.1* in citrus leaves ^[26]^. Thus, *CsGH3.1* and *CsGH3.1L* appear to be functional IAA-amido synthetase genes involved in the regulation of free IAA levels in citrus. *CsGH3.1* and *CsGH3.1L* belong to the group II proteins of the GH3 family. *Arabidopsis GH3.5* and rice *GH3.1* and *sGH3.8* of this group positively regulate pathogen-induced defense responses through the depletion of free IAA ^[13, 24, 25]^. Similarly, the overexpression of *CsGH3.1* and *CsGH3.1L* enhances plant resistance to citrus canker. Moreover, we showed that the overexpression of *CsGH3.1* and *CsGH3.1L* accelerated the Xcc-induced decline of free IAA levels in transgenic plants. Based on these data, our results indicated that the overexpression of *CsGH3.1* and *CsGH3.1L* reduces the susceptibility to *Xcc* by decreasing free IAA levels both before and after pathogen infection in citrus.

The transcriptomic data showed that the decrease in the free IAA level significantly repressed the expression levels of auxin signaling-related genes in transgenic plants. For example, AUX/IAA family members, SAUR-like auxin-responsive protein family members, and ARF10, PIN1, PIN3 and AUX1-like protein genes, which are involved in auxin signal transduction, were significantly downregulated in transgenic plants. In particular, the expression levels of the AUX/IAA family members annotated by our transcriptomic data were significantly repressed by the overexpression of CsGH3. The AUX/IAA family encodes a group of primary auxin-responsive proteins, which regulate ARF expression through ubiquitin-mediated degradation ^[17]^. When cellular auxin concentrations are below a certain threshold, AUX/IAA proteins inhibit ARF transcription factors to activate auxin responsive genes (such as *AUX/IAA, GH3* and *SAUR*) and subsequently suppress auxin responses ^[9]^. This suppression further represses AUX/IAA transcript levels. The decreased expression of PIN1 and PIN3 indicated that the auxin efflux was inhibited in transgenic plants, suggesting that the overexpression of *CsGH3.1* and *CsGH3.1L* affected the auxin distribution in plants. The transcriptomic data also showed that no auxin biosynthesis-related genes (such as those related to tryptophan biosynthesis or metabolism) were affected by the decrease in the free IAA content, while the two genes encoding UDP-glycosyltransferase (UGT74B1 and UGT75B1), which conjugate auxin to glucoside ^[38]^, displayed significantly increased expression levels in transgenic plants. Thus, the unchanged expression levels of auxin biosynthesis-related genes and activation of auxin glucosylation-related genes further favored decreases in the free IAA contents of transgenic plants. Additionally, the IAA–amino acid conjugated hydrolase genes (Cs3g19760 and Cs7g08080), which can rapidly regenerate IAA from IAA–amino acid conjugates to help maintain auxin homeostasis ^[39]^, displayed increased expression levels in the L-5 line, which could be an antagonistic response to depleted free auxin levels.

The constitutive expression of *CsGH3.1* and *CsGH3.1L* affected the establishment of citrus’ architecture. Abnormal phenotypes also occurred in transgenic rice overexpressing *OsGH3.1* ^[25]^ and *OsGH3.8* ^[24]^. The GO annotation revealed that many of the DEGs in transgenic lines were classified into metabolic process, cellular process and development. Cell walls play critical roles in establishing plant architecture. In our study, most genes that were involved in cell wall biosynthesis, were profoundly upregulated by the overexpression of *CsGH3.1* and *CsGH3.1L*. Additionally, three genes (pectinesterase inhibitor, expansin and polygalacturonase), related to cell wall loosening, had decreased expression levels. However, Domingo et al. ^[25]^ showed that most of the genes involved in both cell wall biosynthesis and loosening were downregulated by OsGH3.1’s overexpression in rice. This transcriptional differences of these genes between the two studies indicated that GH3.1 has various functions in the regulation of cell wall-related genes in different species. However, it is clear that overexpressing GH3.1 can result in similar dwarf phenotypes in both rice and citrus.

Auxin is believed to act as a pathogenic factor in pustule formation during citrus canker development ^[5, 16, 26]^. The transcriptional activator-like effectors, PthAs, secreted by Xcc manipulate multiple disease susceptibility genes or their products to regulate pustule formation in infected sites ^[40]^. For example, the PthA-induced expression of the susceptibility genes *CsLOB1, CsLOB2* and *CsDiox* regulate citrus cell division and growth to increase pustule formation ^[3, 4, 40]^. PthA effectors also interact with CsCYP and CsMAF1 proteins, which are repressors of citrus RNA polymerase (Pol) II and III, respectively, to activate the transcription of host genes involved in cell division and growth ^[41]^. Auxin inhibits the translocation of CsMAF1 from the nucleoplasm to nucleolus ^[42]^, which decreased the accumulation of CsMAF1 in the nucleolus. This decreased CsMAF1 content in the nucleolus is beneficial for Pol III’s activation of host genes’ transcription as well as for the PthA effectors’ enhancement of *CsLOB1, CsLOB2* and *CsDiox* expression levels, which both increase symptom development. Thus, conversely, this depletion of free IAA in transgenic plants is favorable for CsMAF1’s entry into the nucleolus to antagonize Pol III-mediated gene expression and finally represses disease development. Moreover, *CsGH3*-mediated auxin homeostasis probably affects *CsLOB1* functions in citrus canker development. The cell wall is the first line of plant defense against pathogen invasion and pathogen-induced host cell wall loosening plays an important role in symptom development ^[43, 44]^. CsLOB1 upregulates the expression of pectate lyase, extension, α-expansin and cellulose genes ^[3]^, which are involved in the cell wall loosening induced by pathogen infection ^[1]^. Auxin triggers cell wall loosening by rapidly acidifying cell walls ^[45]^. Thus, pathogen-induced increases in auxin may have synergistic roles in the cell wall loosening induced by CsLOB1. Our transcriptomic data also showed that a decrease in free auxin significantly repressed the expression levels of cell wall loosening-related genes. Similarly, Cernadas and Benedetti ^[5]^ showed that the auxin transport inhibitor naphthyl-phthalamic acid repressed pustule formation and the expression of cell wall loosening-related genes induced by Xcc infection in the sweet orange (*C. sinensis*) “Pêra” cultivar. Thus, the data showed that interfering with auxin homeostasis, as seen with CsGH3, can weaken pathogen effector-induced host pustule formation and finally enhance plant resistance.

The MapMan analysis showed that biotic stress-related pathways were significantly upregulated by the overexpression of *CsGH3.1* and *CsGH3.1L*. In addition, a considerable number of induced genes were disease-resistance response genes, such as TIR, PR, LRR and TIR-NBS-LRR family members, suggesting that the depletion of the IAA content enhanced the defense response. In the complex network of regulatory interactions during plant resistance responses, an antagonistic relationship between SA and JA signaling pathways is evident, and generally, plant responses to bacterial infections involve activating SA signaling, which represses JA signaling ^[46, 47]^. In citrus, SA treatments enhance resistance to citrus canker ^[48]^. Moreover, SA also inhibits pathogen spread in plants by repressing auxin signaling ^[14]^, and *Arabidopsis GH3.5* enhances the SA-mediated defense response ^[49]^. Thus, we evaluated the effects of overexpressing *CsGH3.1* and *CsGH3.1L* on the hormone contents of transgenic citrus. The transgenic lines had significantly increased SA levels. Additionally, the transcriptomic analysis showed that three of four genes encoding S-adenosyl-L-methionine-dependent methyltransferase superfamily proteins, which convert active SA to non-active methylSA, were repressed in transgenic lines, which was consistent with increased levels of SA. This indicated that the depletion of the IAA content enhanced the SA accumulation by regulating the conversion of SA and methylSA. In JA signaling, the 12-oxophytodienoic acid reductase and cystathionine Beta-synthase genes participating in JA biosynthesis were downregulated in the 1-9 line, which was consistent with the decreased JA level in this line. However, the JA content was not significantly different compared with WT plants, although the two genes were induced in the L-5 line. Thus, our data showed that the overexpression of *CsGH3.1* and *CsGH3.1L* enhanced SA signaling and partially repressed JA signaling, which may activate the expression of disease resistance genes in transgenic plants. The data also showed that ET levels were significantly induced by the overexpression of *CsGH3.1* in transgenic lines. Plants produce ET in response to most biotic and abiotic stresses. In some cases, the role of ET in plant defense contributes to pathogen resistance. In *A. thaliana*, SA and ET function together to coordinately induce several defense-related genes, and ET treatments potentiate the SA-mediated induction of PR-1 ^[47, 50]^. However, the ET signaling pathway may also negatively affect SA-dependent resistance ^[46, 51, 52]^. In our study, *CsGH3.1* and *CsGH3.1L* positively regulated SA and ET accumulations in citrus, which should improve resistance to citrus disease.

In this study, transgenic plants overexpressing *CsGH3.1* and *CsGH3.1L* displayed a similar altered morphology, decreased free IAA levels and enhanced citrus canker resistance. However, a transcriptomic analysis showed that there were obvious differences in the affected MapMan pathways between lines 1-9 and L-5, indicating that *CsGH3.1* and *CsGH3.1L* have different roles in the regulation of auxin signaling. Based on the findings, we hypothesized that *CsGH3.1* and *CsGH3.1L* can increase resistance against citrus canker in citrus plants by inhibiting the accumulation of active auxin, revealing a potential role for the *GH3* gene in citrus breeding to improve citrus canker resistance.

## Author contribution statement

XZ designed the experiments and wrote the manuscript. LJ performed the transcriptomic sequencing and analysis. ZK performed the hormone content analysis. PA performed and evaluated the resistance to citrus canker. CM performed the citrus genetic transformations. LQ performed the phenotype analysis of transgenic plants. HY performed the PCR analyses. SC analyzed the data. All the authors read and approved the manuscript.

## Acknowledgements

This work was supported by the National Key Research and Development Program of China (2018YFD1000300), the Natural Science Foundation Project of CQ (cstc2017jcyjBX0020), the Earmarked Fund for China Agriculture Research System (CARS-27 to S. Chen) and the Science and Technology Major Project of Guangxi (Gui Ke AA18118046). We thank Lesley Benyon, PhD, from Liwen Bianji, Edanz Group China (www.liwenbianji.cn/ac), for editing the English text of a draft of this manuscript.

